# Perceived multisensory common cause relations shape the ventriloquism effect but only marginally the trial-wise aftereffect

**DOI:** 10.1101/2024.08.12.607537

**Authors:** Christoph Kayser, Herbert Heuer

**Author notes:** Corresponding author: Christoph Kayser.

## Abstract

Combining multisensory cues is fundamental for perception and action, and reflected by two frequently-studied phenomena: multisensory integration and sensory recalibration. In the context of audio-visual spatial signals, these are exemplified by the ventriloquism effect and its aftereffect. The ventriloquism effect occurs when the perceived location of a sound is biased by a concurrent visual stimulus, while the aftereffect manifests as a recalibration of sound localization after exposure to spatially discrepant stimuli. The relationship between these processes—whether recalibration is a direct consequence of integration or operates independently—remains debated. This study investigates the role of causal inference in these processes by examining whether trial-wise judgments of audio-visual stimuli as originating from a common cause influence both the ventriloquism effect and the immediate aftereffect. Using a spatial paradigm, participants made explicit judgments about the common cause of stimulus pairs, and their influence on both perceptual biases was assessed. Our results indicate that while multisensory integration is contingent on common cause judgments, the immediate recalibration effect is not. This suggests that recalibration can occur independently of the perceived commonality of the multisensory stimuli, challenging the notion that recalibration is solely a byproduct of integration.

## Introduction

Combining multiple sensory cues is central to perception and action during daily activities. Current research differentiates two kinds of such multisensory combinations: the integration of discrepant signals, as revealed by perceptual judgments of multisensory stimuli, and the recalibration of one or both signals, as revealed by the aftereffect of discrepant stimuli on subsequent judgments of unisensory stimuli. In the context of audio-visual spatial signals, these multisensory combinations are exemplified by the ventriloquism effect and aftereffect (Bertelson & Radeau, 1981; Recanzone, 1998). The ventriloquism effect reflects how our judgements of (e.g.) a sound’s location are biased by seeing a visual stimulus at the same time. The aftereffect reflects how exposure to spatially discrepant audio-visual stimuli biases the localization of subsequently presented sounds and grows cumulatively with the exposure to discrepant stimuli. The relation between the ventriloquism effect and aftereffect, and thus the underlying processes of integration and recalibration, is still debated.

One line of studies supports that the aftereffect is a direct consequence of the preceding integration of multisensory signals (Bruns, 2019; Rohlf *et al*., 2020; Noppeney, 2021; Rohlf *et al*., 2021). Both integration and recalibration scale similarly with the discrepancy of the multisensory stimuli, and this discrepancy may directly drive both processes (Wozny & Shams, 2011b; Bruns & Roder, 2015; Park & Kayser, 2019). Furthermore, both are similarly affected by manipulations of stimulus reliability (Rohlf *et al*., 2021) and attention (Badde *et al*., 2020) and neuroimaging studies point to partly overlapping neurophysiological processes shaping both (Park & Kayser, 2019; 2021). It is known that attributing a common cause to multisensory stimuli is prerequisite to their features being integrated (Ernst & Bulthoff, 2004; Wallace *et al*., 2004; Kording *et al*., 2007; Rohe & Noppeney, 2015). Indeed, the perceptual bias arising from integration emerges only in trials in which the two stimuli are judged as being causally related. If recalibration was a direct consequence of integration, recalibration should also depend on whether the two spatially discrepant stimuli are judged as originating from a common source. However, it remains unclear whether this is indeed the case.

Another line of studies argues that recalibration is driven mainly by a belief in a modality-specific bias. In the case of the ventriloquism aftereffect, this would be a bias which is specific to estimates of acoustic features that should emerge independently of whether the preceding multisensory signals have been integrated or not (Di Luca *et al*., 2009; Block & Bastian, 2011; Zaidel *et al*., 2013; Bosen *et al*., 2017). If this notion was correct, recalibration would emerge independently of whether the multisensory stimuli are judged as being causally related or originating from a common source.

These predictions should be particularly true on a trial-by-trial level. Recalibration is observed following a single trial of audio-visual exposure (known as immediate or trial-wise aftereffect), but also growths cumulatively with extended exposure. Hence, recalibration depends on both the sensory information received during an individual trial as well as the consistency of this over an extended period (Bruns & Roder, 2015; Bosen *et al*., 2017). Indeed, when probed across sequences of multisensory trials integration depends only on the currently presented spatial discrepancy while recalibration is sensitive to the series of preceding spatial discrepancies (Kayser *et al*., 2023). The different time scales on which integration and recalibration operate could come along with different time scales of dependence on multisensory common cause evidence (Debats *et al*., 2023a). Thus, whereas on a given trial multisensory integration emerges only for a pair of discrepant multisensory stimuli attributed to a common cause in that particular trial, immediate recalibration following these stimuli may only minimally depend on this short-term common cause evidence.

The present study was designed to arbitrate between these alternatives by probing the influence of trial-wise common cause judgements on the ventriloquism effect and the immediate aftereffect in an audio-visual spatial paradigm. This paradigm has been used previously to study the ventriloquism effect and the immediate (i.e. trial-wise) aftereffect in different contexts (Park & Kayser, 2019; 2020; 2022; Kayser *et al*., 2023). We here expanded this paradigm by including explicit trial-wise judgements about participants believe that audio-visual stimulus pairs were originating from a common cause or not, and tested the predictive power of these common cause judgements on both biases. Our results show that the integration of discrepant stimuli depends on whether they are judged as originating from a common cause, as expected, while the immediate ventriloquism aftereffect does not.

## Methods

### Participants

We report data from two experiments in which 44 adult volunteers participated after providing informed consent. All had self-reported normal vision and hearing and none indicated a history of neurological disorders. The procedures were approved by the ethics committee of Bielefeld University and the data were collected anonymously. Participants were compensated financially for their time. For experiment 1 we collected data from 23 participants, for experiment 2 from 21 participants. Experiment 2 was implemented to reproduce the results obtained in experiment 1 using a different type of common cause response method, as explained below. These sample sizes are in line with previous studies using similar experimental protocols (Park & Kayser, 2019; Park *et al*., 2021; Park & Kayser, 2022) and recommendations for behavioral studies (Simmons *et al*., 2011).

### Experimental setup and paradigm

The experiments were based on an established single-trial audio-visual localization task in which the individual trials either probe the ventriloquism effect or the immediate ventriloquism aftereffect (Wozny & Shams, 2011b; Park & Kayser, 2019; 2021; 2022). They were conducted in an echo-free booth (E:Box, Desone) where participants sat in front of an acoustically transparent screen (Screen International Modigliani, 2×1 m^2^, about 1m from the participants head) with their head supported on a chin rest. The experiments comprised audio-visual (AV) trials and auditory (A) trials, and less frequent visual (V) trials. These visual trials were mainly included to maintain attention divided across modalities. Participants’ task was to localize either the sound (in AV or A trials) or the visual stimulus (in V trials). They were asked to fixate a central dot during the pre-stimulus and stimulus periods, but could move their eyes during responding and inter-trial intervals.

Each trial started with a fixation period (uniform 700 to 1100 ms), followed by stimulus presentation (50 ms). After a post-stimulus interval (uniform 400 to 700 ms) participants had to indicate the position of the auditory (in AV and A trials) or visual (in V trials) stimulus. A horizontal bar was presented along which participants could move a mouse cursor to indicate their judgement by pressing the mouse button. In AV trials, participants were subsequently asked (after a uniform interval of 200 to 300 ms) to indicate whether they perceived the auditory and visual stimuli as ‘to arise from the same source’ or ‘from different sources’ (common cause judgement). In experiment 1 participants were instructed to move a mouse cursor along a vertical bar to a position that would mark the strength of their belief about a common cause; the two extreme positions were labeled ‘common cause’ (highest position) and ‘separate causes’ (lowest position).

After running experiment 1 we realized that several participants tended to move the mouse cursor to the extreme positions and effectively submitted binary decisions. For this reason, we implemented experiment 2, in which the common cause judgement was directly a binary decision, with movements of the mouse cursor upwards from the midline labelled as ‘common cause’ and those downwards as ‘separate causes’. In both experiments’ participants finalized the common cause judgment by clicking a mouse button. Trials were separated by inter-trial intervals between 800 and 1200 ms.

Stimulus presentation was controlled using the Psychophysics toolbox (Brainard, 1997) for MATLAB (The MathWorks Inc., Natick, MA) with confirmed temporal synchronization of auditory and visual stimuli. The acoustic stimuli were presented through speakers (Monacor MKS-26/SW, MONACOR International GmbH & Co. KG) that were located behind the visual screen at 5 discrete horizontal locations (±22°,±11°, 0°), with the center position always being directly in front of the participant. Sound presentation was controlled via a multi-channel soundcard (Creative Sound Blaster Z) and amplified via an audio amplifier (t.amp E4-130, Thomann). The acoustic stimulus was a 1300 Hz sine wave tone (50 ms duration; sampled at 48 kHz) that was presented at one of two signal-to-noise ratios. These were introduced to manipulate the reliability of the acoustic spatial information and thereby the integration bias and possibly the common cause judgments. In the high reliability condition (labelled A+) the tone at the target position was presented at 65 dB SPL, while the same tone was presented from all other four speakers at ∼45 dB SPL. In the low reliability condition (labelled A-) the target was presented at 58 dB, hence with less intensity difference to the other speakers.

Visual stimuli were clouds of dots distributed according to a two-dimensional Gaussian distribution (200 dots, SD of vertical and horizontal spread = 2°, width of a single dot = 0.12°, duration 50 ms) similar to previous studies (Park & Kayser, 2019; 2021). They were projected (Acer Predator Z650, Acer Inc., Taiwan) onto the acoustically transparent screen. Auditory and visual stimuli were presented at the same or different discrete locations, inducing a multisensory spatial discrepancy of various sizes in the majority of AV trials. The specific locations of auditory and visual stimuli within trials (for AV trials) and between trials were drawn semi-independently.

Experiment 1 featured a total of 832 trials administered in 6 blocks (of these 360 AV trials, 360 A trials, and 112 visual trials). The audio-visual discrepancies ranged from -44° to +44° in steps of 11°. The largest discrepancies of ±44° were presented in only a few AV trials, and the responses proved rather variable. Therefore this discrepancy was not included in the design of experiment 2, which featured a total of 742 trials administered in 4 blocks (of these 306 AV trials 306, A trials and 126 visual trials), with discrepancies ranging from -33° to + 33°.

One or two blocks of practice trials were run prior to the main experiment to familiarize participants with the task and stimuli. During these trials feedback was provided by dividing the judgement error into three classes and providing respective feedback (‘good’, ‘intermediate’, ‘off’). Before the actual experiment participants were also screened for their spatial hearing abilities. They were asked to localize noise bursts (50 ms duration, 65 dB SPL) presented from the 4 lateralized speakers in a left/right two-choice task, performing 10 repeats per location. All participants performed above an average of 75% correct responses.

### Definition of trial-wise judgement errors

The trial-wise judgement errors were defined so as to take potential response biases into account, such as a bias towards the midline of the screen (Kording *et al*., 2007; Wozny & Shams, 2011a; b; Park & Kayser, 2019). Practically, we computed the deviation between the participant’s trial-wise response and the expected response given the location of the judged sound in that trial, similar to previous work. This expected response was computed for each sound location as the average response across all trials with a sound at this location. In line with previous studies (Kayser *et al*., 2023; Kayser *et al*., 2024) we removed individual trials with judgement errors larger than 150% of the largest spatial discrepancy included in each experiment. These trials can reflect lapses of attention or accidental clicks when submitting the response. From experiment 1 we also removed trials in which the spatial discrepancy was ±44°, as these responses exhibited a much larger trial-to-trial variability than those at other discrepancies.

### Analysis of common cause judgements

In experiment 1 common cause judgements were provided on a continuous scale, but participants often used only the extremes of the continuum. Hence, we converted the common cause judgements to a binary variable for all participants. To account for potential individual biases in how participants used the continuous scale, we split the judgements of each participant by the overall arithmetic mean judgement for this participant. Note that we used the mean and not the median to split trials, as for some participants the median was one of the two extreme values. Still, for 4 of the 23 participants this left not sufficient variability in the common cause judgments and these participants were excluded. In experiment 2 participants’ responses were directly saved as a binary response for each trial and were used as such for analysis. Comparing the common cause judgements across experiments confirms that these are very similar (Fig. 1 and 2), and overall the results from both experiments conform very well.

**Figure 1.**
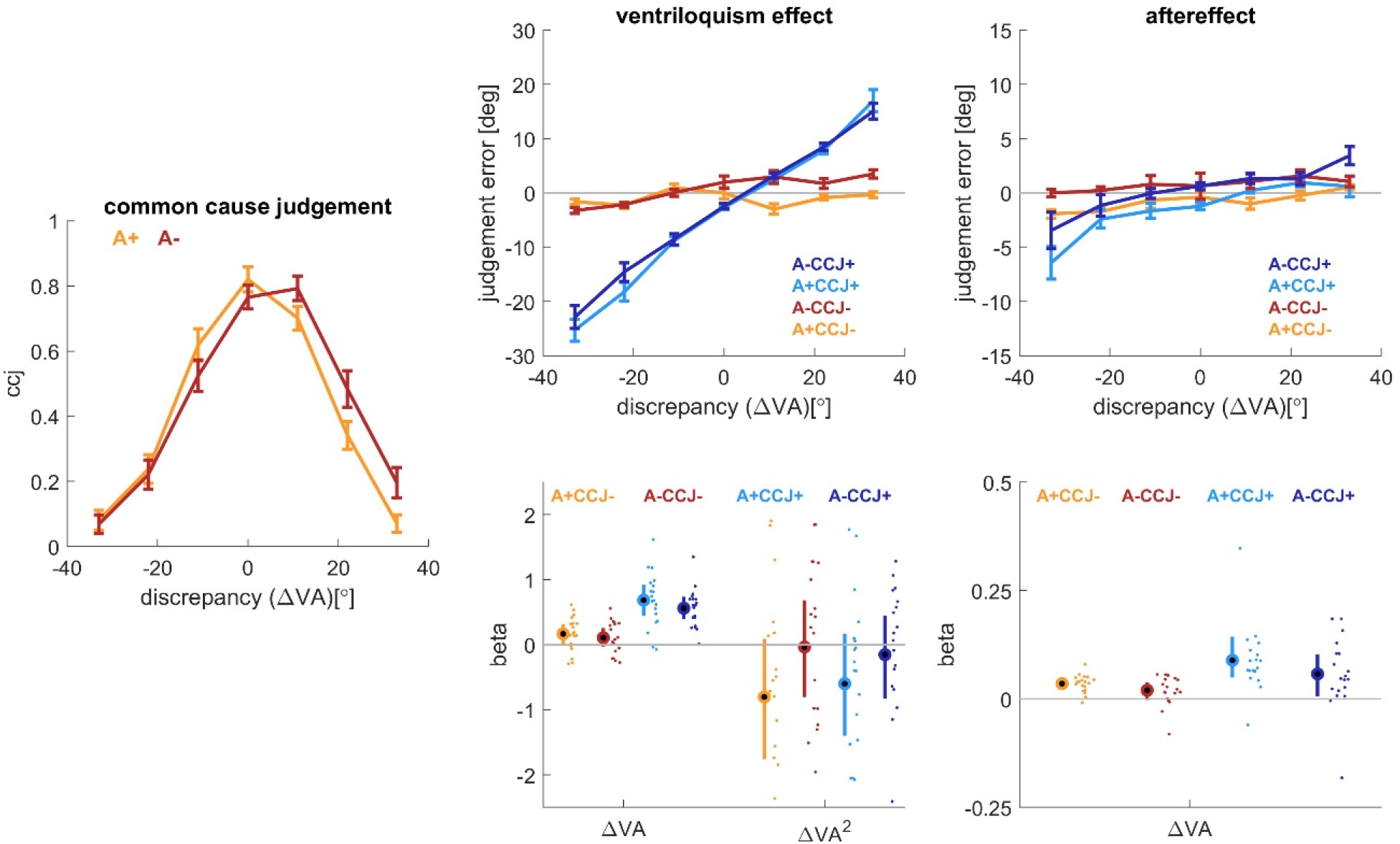
Results for experiment 1 (n=19). Left panel: Common cause judgments as a function of the spatial discrepancy and auditory reliability (high: A+, low: A-). Middle panel: Ventriloquism effect, Right panel: Aftereffect. Both are shown as the respective continuous judgement errors for each of the conditions (upper panel) and as participant-wise slopes against the spatial discrepancy (lower panel). These individual slopes were derived by fitting the models to the individual participant data and are mainly intended to show the effects of interest and their variability across participants. Lines indicate means and standard errors, dots individual participants. CCJ+: trials on which participants rated the audio-visual stimuli as to arise from a common cause; CCJ-: trials on which participants rated them as to arise from separate sources. Note the different scales on the ordinates for ventriloquism effect and aftereffect.

**Figure 2.**
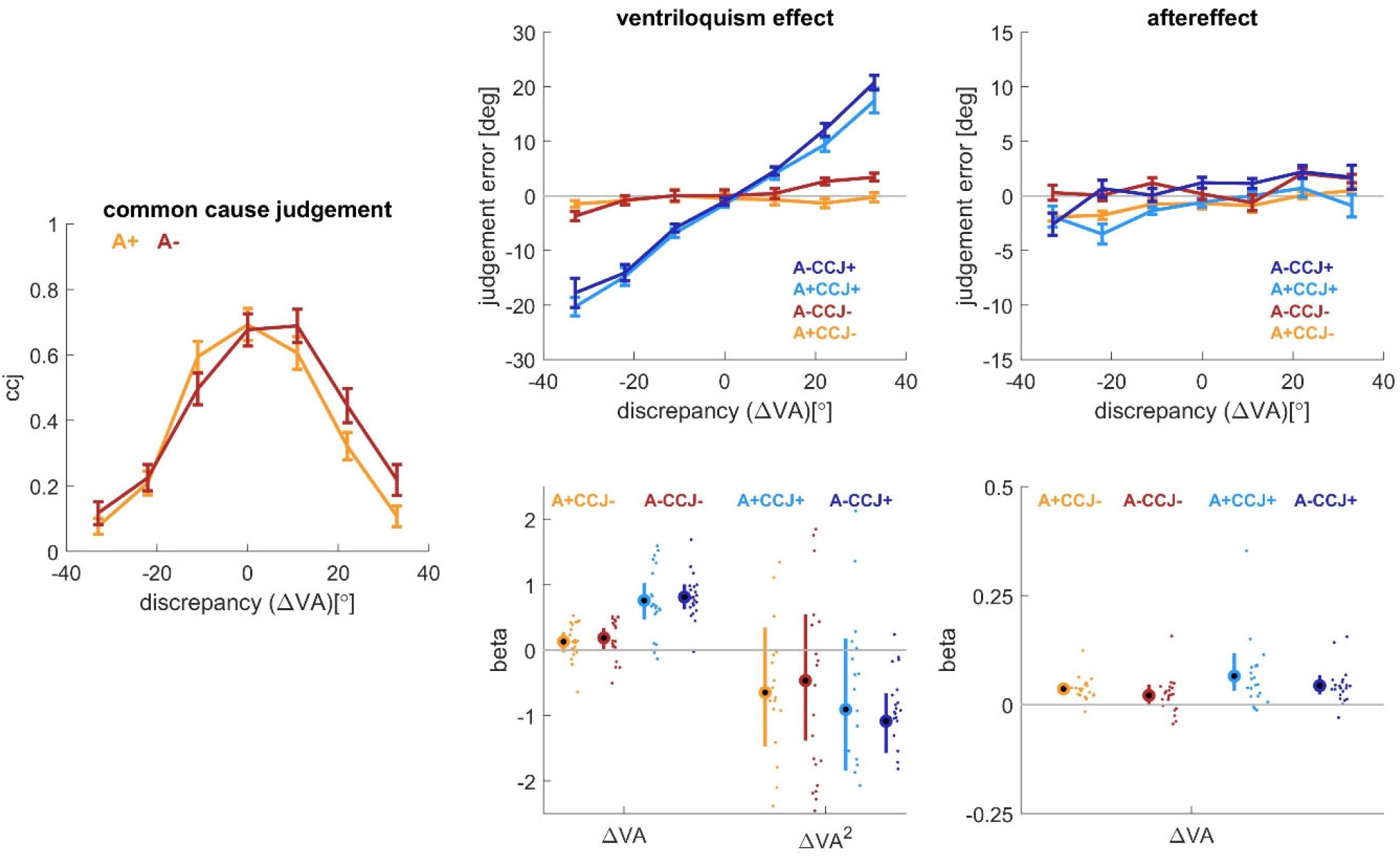
Results for experiment 2 (n=21). Left panel: Common cause judgments as a function of the spatial discrepancy and auditory reliability (high: A+, low: A-). Middle panel: Ventriloquism effect, Right panel: Aftereffect. Both are shown as the respective continuous judgement errors for each of the conditions (upper panel) and as participant-wise slopes against the spatial discrepancy (lower panel). These individual slopes were derived by fitting the models to the individual participant data and are mainly intended to show the effects of interest and their variability across participants. Lines indicate means and standard errors, dots individual participants. CCJ+: trials on which participants rated the audio-visual stimuli as to arise from a common cause; CCJ-: trials on which participants rated them as to arise from separate sources. Note the different scales on the ordinates for ventriloquism effect and aftereffect.

### Analysis of the ventriloquism bias and aftereffect

The ventriloquism bias was defined based on the judgment errors in the AV trials, the aftereffect was defined based on the judgment errors in the A trials. These judgement errors scale with the audio-visual spatial discrepancy ΔVA, defined as the location of the visual stimulus minus that of the sound in the AV trials. This scaling can comprise both linear and non-linear patterns, in particular for the ventriloquism bias. The non-linear dependency describes the reduced tendency to bind multisensory stimuli when these are seemingly discrepant and not judged as arising from a common cause (Kording *et al*., 2007; Rohe & Noppeney, 2015; Shams & Beierholm, 2022), following Bayesian models of sensory causal inference. Similar to previous work we modeled a nonlinear dependency as the signed square-root of the magnitude of the discrepancy (*ΔVA*^*2*^ *= sign(*ΔVA*) * sqrt(*ΔVA)) (Park & Kayser, 2020; 2022). In a first analysis we tested whether the data for each bias are better accounted for by a linear or a combined linear and nonlinear model. We then used the best-fitting model to test for the effects of interest: the modulation of the dependency of the biases on ΔVA by stimulus reliability and the trial-wise common cause judgment.

### Statistical analysis and hypothesis testing

Statistical analyses were based on generalized linear models, which were fit separately to the trial-wise judgment errors for the ventriloquism effect or the aftereffect for each experiment. In a first step we probed whether each bias is best described by linear (ΔVA) and or nonlinear (*ΔVA*^*2*^) dependencies on the spatial discrepancy. For this we compared the predictive power of three models:

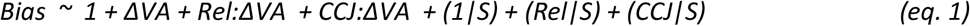

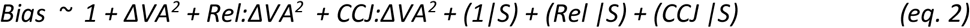

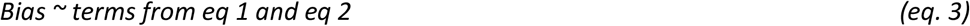

Here S denotes the participant, *Rel* the reliability (two levels) and *CCJ* the common cause judgment (two levels). Models were fit using a maximum likelihood procedure using the Laplace method in Matlab R2021a (fitglme.m). Models were compared based on their respective BIC values. This revealed that for both experiments the ventriloquism effect was best accounted for by the combined linear and nonlinear model (delta BIC relative the second-best model Exp 1: 16.9; Exp2: 15.8), while the aftereffect was best accounted for by a linear model (delta BIC: 5.1 and 6.6). We hence relied on the combined model (eq. 3) for the ventriloquism effect and the linear model (eq. 1) for the aftereffect, similar to previous work.

When testing for the effects of interest (reliability and common cause judgment) we relied on the differences in BIC values between the complete model including the predictor of interest (eq. 3 for the ventriloquism effect, eq. 1 for the aftereffect) and the reduced model omitting this particular factor (e.g. CCJ); note that omitting a predictor refers to both the interaction with the ΔVA terms and the random effects. Differences in BIC values were converted to Bayes factors. With this we base our statistical conclusions not on the significance of individual model coefficients but rather emphasize the predictive power of individual factors in the model; we also base our conclusions on Bayes factors rather than classical null hypothesis tests in line with recommendations for behavioral studies to use Bayesian statistics (Wagenmakers, 2007; Wagenmakers *et al*., 2018). Bayes factors in favor no effect of a predictor are reported as negative numbers, rather than as fractions smaller than one (i.e. Bayes factors smaller than 1 were converted as -1/BF). When interpreting Bayes factors, we refer to the nomenclature of Raftery (Raftery, 1995) and interpret BF between 1 and 3 as ‘weak’, between 3 and 20 as ‘positive’, between 20 and 150 as ‘strong’, and BF > 150 as ‘very strong’ evidence. The model coefficients are reported in Tables 2 and 3. To quantify the contribution of the CCJ to the ventriloquist effect and the aftereffect we also calculated the variance (adjusted R^2^) explained by each model and percentage of variance lost when ignoring the CCJ as factor; this was defined as 100*(R^2^ full model – R^2^ reduced model) / R^2^ full model. To visualize the effects of auditory reliability and common cause judgements we calculated the regression models for individual participants. The individual participant slopes of each factor are shown in Figures 1 and 2 bottom panels.

For the analysis of the common cause judgements we relied on a similar model, predicting the trial-wise (binary) common cause judgement based on the reliability, the magnitude of the discrepancy and their interaction (using a binomial model with a logistic link function). Again we compared the BIC values of models including or omitting a factor of interest. Overall we report results for 19 participants in experiment 1, based on 275±16 AV/A trial-pairs (mean±SD) and results for 21 participants in experiment 2, based on 303±13 AV/A trial-pairs.

## Results

### Common cause judgements

The common cause judgements (CCJ) scaled with the audio-visual discrepancy, and participants judged two stimuli as less likely to emerge from a common cause when separated by a larger distance (Fig 1 and 2, left panels). For statistical analysis we modelled the trial-wise CCJ against the magnitude of the spatial discrepancy, the reliability of the auditory stimuli, and their interaction (Table 1); we quantified the predictive power of either the reliability or the discrepancy by contrasting models including or omitting this factor of interest. For both experiments this revealed very strong evidence in favor of a role of the spatial discrepancy in predicting the CCJ (delta BIC: 1694 and 1398 for experiment 1 and 2; corresponding BF ∼10^10^) and no clear evidence in favor of a role of the auditory reliability (delta BIC: -22 and 3; BF ∼-10^5^ and 5.6). This strong dependency of the CCJ on the degree of spatial discrepancy confirms previous studies using a similar paradigm (Rohe & Noppeney, 2015).

**Table 1.**
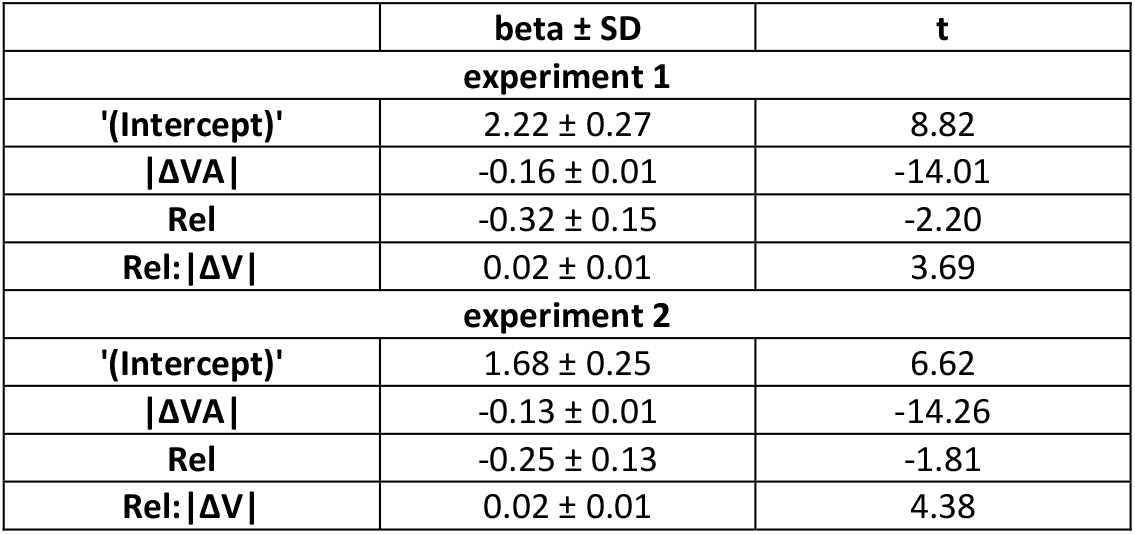
Influence of auditory reliability and multisensory discrepancy on common cause judgements. For each experiment we modelled the trial-wise common cause judgment against the magnitude of the spatial discrepancy (|*Δ*va|), the reliability of the acoustic signal (levels high reliability, low reliability) and their interaction. The table lists the respective coefficients of each factor including their standard deviation and the respective t-values.

### Ventriloquism effect

As expected based on previous work, the judgement errors reflecting the ventriloquism bias scaled with the spatial discrepancy both in a linear and nonlinear manner (see Methods) and were shaped by the reliability of the acoustic stimuli (Table 2). Comparing models including or omitting the factors of interest provided very strong evidence for a role of reliability in predicting the ventriloquism bias (delta BIC: 57 and 124 for experiments 1 and 2; both BF ∼10^10^) and very strong evidence for a role of the common cause judgements in doing so (delta BIC: 2183 and 2097; both BF ∼10^10^). This influence of the common cause judgment on the ventriloquism effect was particularly pronounced at larger spatial discrepancies (Figure 1 and 2, middle panels) and was consistent across participants (lower panels). To provide an additional measure of the contribution of the common cause judgement to the ventriloquism effect, we calculated the percentage of explained variance lost when removing this predictor: this amounted to 53% and 56% of lost variance for experiments 1 and 2 respectively.

**Table 2.** Influence of auditory reliability and common cause judgements on the ventriloquism effect. For each experiment we modelled the trial-wise judgement errors against the linear spatial discrepancy (*Δ*va), a nonlinear dependency on discrepancy (*Δ*va^2^), and the interactions of these with the auditory reliability (Rel) and the common cause judgement (CCJ) as explained in the Methods (eq. 3).

| | beta ± SD | t |
| --- | --- | --- |
| **Experiment 1** |  |  |
| '(Intercept)' | -0.69 ± 0.12 | -5.44 |
| $\Delta VA$ | -0.50 ± 0.09 | -5.22 |
| $\Delta VA^2$ | -0.39 ± 0.09 | -0.81 |
| Rel:$\Delta VA$ | 0.04 ± 0.04 | 1.02 |
| CCJ: $\Delta VA$ | 0.60 ± 0.05 | 11.98 |
| Rel: $\Delta VA^2$ | -0.5 ± 0.21 | -0.22 |
| CCJ: $\Delta VA^2$ | -0.29 ± 0.23 | -1.23 |
| **Experiment 2** |  |  |
| '(Intercept)' | -0.17 ± 0.17 | -0.97 |
| $\Delta VA$ | -0.66 ± 0.09 | -6.73 |
| $\Delta VA^2$ | 0.37 ± 0.50 | 0.75 |
| Rel:$\Delta VA$ | 0.14 ± 0.04 | 3.31 |
| CCJ: $\Delta VA$ | 0.60 ± 0.05 | 11.79 |
| Rel: $\Delta VA^2$ | -0.41 ± 0.22 | -1.83 |
| CCJ: $\Delta VA^2$ | -0.34 ± 0.24 | -1.41 |

### Aftereffect

The main focus of this study was the influence of the common cause judgement on the aftereffect. Figure 1 and 2, right panels, show the aftereffect, as the respective judgement errors and as slopes of this judgement error against the spatial discrepancy. As known from previous work (Park & Kayser, 2021; 2022), the slopes of the aftereffect are smaller than those of the ventriloquism effect, but were consistently positive.

For both experiments did we obtain very strong evidence for a role of reliability in predicting the aftereffect (delta BIC 89 and 63; both BF ∼10^10^; Table 3 for the individual model coefficients). Most importantly, for both experiments did we obtain very strong evidence against a role of the common cause judgements in predicting the aftereffect (delta BIC: -22 and -31; BF=-1177 and -10^7^). Furthermore, the percentage of variance lost in the explanatory power for the aftereffect when ignoring the common cause judgement was 7.8% and 1.4% respectively, and hence much smaller compared to loss for the ventriloquism effect (above 50%). All in all this shows that the common cause judgement has little predictive power for the aftereffect.

**Table 3.** Influence of auditory reliability and common cause judgements on the aftereffect. For each experiment we modelled the trial-wise judgement errors against the linear spatial discrepancy (*Δ*va) and the interactions of this with the auditory reliability (Rel) and the common cause judgement (CCJ) as explained in the Methods (eq. 1).

| | beta ± SD | t |
| --- | --- | --- |
| **Experiment 1** |  |  |
| '(Intercept)' | -0.082 ± 0.083 | -0.98 |
| $\Delta VA$ | 0.007 ± 0.015 | 0.47 |
| Rel:$\Delta VA$ | -0.014 ± 0.007 | -1.95 |
| CCJ: $\Delta VA$ | 0.042 ± 0.009 | 4.50 |
| **Experiment 2** |  |  |
| '(Intercept)' | 0.062 ± 0.083 | 0.74 |
| $\Delta VA$ | 0.030 ± 0.015 | 2.01 |
| Rel:$\Delta VA$ | -0.014 ± 0.007 | -2.02 |
| CCJ: $\Delta VA$ | 0.018 ± 0.009 | 2.11 |

## Discussion

Multisensory integration and recalibration are two processes by which the brain deals with seemingly discrepant sensory signals. Though both are well investigated, their relation remains a matter of discussion. Multisensory integration supposedly reduces the discrepancy between two redundant sensory signals that are received more or less at the same time and judged as originating from a common cause (Kording *et al*., 2007; Rohe & Noppeney, 2015; Zierul *et al*., 2019; Badde *et al*., 2020; Shams & Beierholm, 2022). Recalibration, in contrast, supposedly serves to reduce persistent or constant discrepancies between sensory estimates, and begins to emerge after a single exposure to discrepant audio-visual stimuli (Bruns & Roder, 2015; Bosen *et al*., 2017; Bosen *et al*., 2018). According to one hypothesis, recalibration is driven by the believe in a modality specific bias, and not strictly tied to the integration of the multisensory stimuli driving recalibration (Block & Bastian, 2011; Zaidel *et al*., 2013; Bosen *et al*., 2017; Bruns & Röder, 2019; Tong *et al*., 2020). We here support this idea and show that the trial-wise belief that two spatially discrepant multisensory stimuli originate from a common cause is a prerequisite for integration but the immediate aftereffect shows little dependence on this. This would explain why the aftereffect can also emerge when auditory and visual signals obviously do not originate from a common location (Radeau & Bertelson, 1974) and when attention is directed towards task-unrelated stimuli (Eramudugolla *et al*., 2011).

### Trial-wise recalibration does not depend on trial-wise common cause judgments

In the type of paradigm used here, the trial-wise aftereffect is statistically correlated with both the spatial discrepancy in the preceding multisensory trial and participants’ trial-wise integration biases (Park & Kayser, 2019; 2021). However, this does not imply that the trial-wise integration causally drives recalibration. In fact, based on the analysis of EEG data we have concluded that the neurophysiological drivers of the trial-wise aftereffect are more shaped by the trial-wise spatial discrepancy rather than the integration bias observed in the multisensory trials (Park & Kayser, 2021). Hence, while both the sensory signals and their resulting interpretation and action in a multisensory trial may feed into recalibration (Park & Kayser, 2021; Park *et al*., 2021), the sensory signals are what directly shapes the trial-wise aftereffect. Based on these results on may predict that while integration is contingent on attributing both stimuli to a common cause, the trial-wise aftereffect should be not; just as observed here.

In line with previous work the current results support the notion that recalibration is shaped by the believe in a modality-specific bias, here one pertaining to the auditory system. Multiple factors may shape this belief, some of which also shape integration; hence both biases often tend to correlate across experimental manipulations. However, these biases show a differential sensitivity to the history of the multisensory signals. When probed across sequences of experimental trials, integration depends only on the discrepancy of the audio-visual signals in the current, while recalibration reflects both the immediate and preceding past (Kayser *et al*., 2023). A similar history dependence may also pertain to the common cause evidence, where integration depends mostly on the momentary common cause evidence while recalibration on this evidence over a longer time. Indeed, in a visuo-motor paradigm we have shown that reducing the inter-trial evidence for a common cause reduces integration but not recalibration (Debats *et al*., 2017; Debats *et al*., 2023b). Hence, recalibration may be more sensitive to the evidence for sensory-causal relations on a longer time scale, while integration is contingent on the immediate common cause evidence.

### Integration and recalibration from a Bayesian perspective

We interpret the collective results obtained previously (Park & Kayser, 2021; 2022; Debats *et al*., 2023b; Kayser *et al*., 2023) as follows: the combination of multisensory signals is generally shaped by models of Bayesian causal inference (Noppeney, 2021; Shams & Beierholm, 2022). Unpredictable discrepancies emerging for stimuli that are assumed to arise from a common cause are reduced on a trial-by-trial level through multisensory integration. At the same time the brain continuously updates the a priori belief about a common cause of the multisensory signals based on the recently experienced discrepancies (Kording *et al*., 2007; Beierholm *et al*., 2020), attention (Badde *et al*., 2020), and the overall context in which the sensory information is received. Separately, at least one of the consistently discrepant sensory signals (e.g. the representation of auditory space here) is continuously updated in a leaky manner to compensate for short-term changes in audio-visual disparities and to minimize apparent localization errors (Kopco *et al*., 2009{Recanzone, 1998 #8206; Wozny & Shams, 2011a; Bosen *et al*., 2018). The estimates of multisensory discrepancies that feed into this update may be indirectly shaped by the common cause evidence, but the data suggest that distinct time scales of this are relevant for integration and recalibration: for integration the immediate common cause evidence is most relevant, while for recalibration it is the long-term evidence – and the immediate evidence may matter only as a potential start of long-term evidence. Dissociating these time scales of common cause is difficult but not impossible, and a manipulation that holds promise to better dissociate the factors shaping integration and recalibration in future work.

## Acknowledgements

This study was funded by the Deutsche Forschungsgemeinschaft (DFG KA2661/2-1). The authors declare that they have no conflicts of interest. We thank Jenny Jakisch and Lisa Stetza for their help during data collection.

## Data availability

The data and relevant Matlab code for producing the figures will be made available under https://github.com/christophckayser

## Abbreviations

BF: Bayes factor

